# A mouse model for spinal muscular atrophy provides insights into non-alcoholic fatty liver disease pathogenesis

**DOI:** 10.1101/2020.04.29.051938

**Authors:** Marc-Olivier Deguise, Chantal Pileggi, Ariane Beauvais, Alexandra Tierney, Lucia Chehade, Yves De Repentigny, Jean Michaud, Maica Llavero-Hurtado, Douglas Lamont, Abdelmadjid Atrih, Thomas M. Wishart, Thomas H. Gillingwater, Bernard L. Schneider, Mary-Ellen Harper, Simon H. Parson, Rashmi Kothary

## Abstract

**Background & aims:** Spinal muscular atrophy (SMA) is an inherited neuromuscular disorder leading to paralysis and death in children. SMA patients are more susceptible to dyslipidemia as well as liver steatosis, features reproduced in SMA mouse models. As current pre-clinical models of NAFLD are invariably imperfect and generally take a long time to develop, the rapid development of liver steatosis in SMA mice provides a means to identify molecular markers of non-alcoholic fatty liver disease (NAFLD). Here, we investigated whether *Smn^2B/-^* mice, a model of severe SMA, display typical features of NAFLD/non-alcoholic steatohepatitis (NASH).

**Methods:** Biochemical, histological, electron microscopy, proteomic, and high-resolution respirometry were used.

**Results:** The *Smn^2B/-^* mice develop steatohepatitis early in life. The consequent liver damage arises from mitochondrial reactive oxygen species production and results in impaired hepatic function including alterations in protein output, complement, coagulation, iron homeostasis, and IGF-1 metabolism. The steatohepatitis is reversible by AAV9-SMN gene therapy. The NAFLD phenotype is likely due to non-esterified fatty acid (NEFA) overload from peripheral lipolysis, subsequent to hyperglucagonemia compounded by reduced muscle use. Mitochondrial β-oxidation contributed to hepatic damage as we observed enhanced hepatic mitochondrial β-oxidation and reactive oxygen species production. Hepatic mitochondrial content, however, was decreased. In contrast to typical NAFLD/NASH, the *Smn^2B/-^* mice lose weight due to their neurological condition, develop hypoglycemia and do not develop hepatic fibrosis.

**Conclusion:** The *Smn^2B/-^* mice represent a good model of microvesicular steatohepatitis. Like other models, it is not representative of the complete NAFLD/NASH spectrum. Nevertheless, it offers a reliable, low-cost, early onset model that is not dependent on diet to identify molecular players in NAFLD pathogenesis and can serve as one of the very few models of microvesicular steatohepatitis for both adult and pediatric populations.

## Introduction

Spinal muscular atrophy is an autosomal recessive disorder characterized primarily by the death of anterior horn motor neurons, leading to paralysis and death. SMA is relatively common with an incidence of 1 in 11,000 live births and a carrier frequency of 1 in 40[1]. The clinical spectrum is wide, spanning severe type I SMA patients, who will never reach major motor milestones such as sitting, and will rapidly succumb to the disease without medical management, to type IV SMA patients with mild weakness[2]. Even though of variable severity, all SMA types arise from a mutation or deletion in the ubiquitously expressed *Survival motor neuron 1* (*SMN1*) gene[3]. The copy number of the *SMN2* gene, a nearly identical gene to *SMN1*, acts as natural genetic modifier and dictates disease severity. The resulting protein from these two genes, SMN, is involved in a number of key cellular pathways, including RNA metabolism and splicing (reviewed in [4]).

SMA has long been defined as primarily a motor neuron disease. Nevertheless, the housekeeping functions of the SMN protein prompt the consideration of defects in non-neuronal cell types. Potential defects in fatty acid metabolism were highlighted in early clinical studies of small cohorts of SMA patients[5–7] but their etiology remains unresolved. More recently, we identified an increased prevalence of dyslipidemia in a cohort of 72 pediatric SMA patients, and increased frequency of fatty liver in comparison to the normal published pediatric population[8]. Strikingly, within the span of two weeks after birth, the *Smn^2B/-^* mouse model of SMA displayed rapid onset of fatty liver disease and dyslipidemia[8]. Apart from this, the *Smn^2B/-^* mouse model shows the typical features of SMA, which include loss of motor neurons, neuromuscular junction abnormalities, skeletal muscle atrophy, muscle weakness, weight loss and a shortened lifespan of 25 days[9]. While incompletely characterized from a metabolic standpoint, *Smn^2B/-^* mice could offer a new model of NAFLD/NASH with a rapid disease onset without the need of long-term diet regimen. As such, these mice could provide a rapid and relatively inexpensive model for new molecular insights in NAFLD/NASH pathogenesis.

NAFLD presents as a spectrum of severity that encompasses simple steatosis, steatohepatitis, cirrhosis and hepatocellular carcinoma[10]. The pathogenesis of NAFLD is complex and involves multiple organ systems. It is currently hypothesized that “multiple hits” are required to develop NASH and more severe phenotypes[10]. Multiple mouse models of NAFLD exist and, as in other diseases, they offer great insight into molecular signaling events. However, these models are invariably imperfect in modelling the true phenotype of patients with NAFLD/NASH[10–13]. In addition, many of these models rely on a dietary component and require a long time to develop the relevant features. Indeed, these two variables can make the current NAFLD/NASH models costly to use.

In a comprehensive analysis of the metabolic defects in *Smn^2B/-^* mice, we show development of NAFLD, and more specifically steatohepatitis without fibrosis, in a very short time span (less than 2 weeks[8]). Of note, the NAFLD was prevented by AAV9-SMN mediated gene therapy. Ultimately, the metabolic defects in *Smn^2B/-^* mice lead to significant functional impairment in key molecular pathways, including general protein production, complement protein expression, coagulation protein expression, insulin-like growth factor 1 (IGF-1) and iron homeostasis pathway regulation. The emergence of the NAFLD phenotype in *Smn^2B/-^* mice is likely from dysfunctional pancreas-liver axis, intrinsic hepatocyte defects and reduced muscle use caused by denervation. Despite showing many features seen in NAFLD, the *Smn^2B/-^* mice do not develop obesity, hyperinsulinemic hyperglycemia, or hepatic fibrosis. Altogether, the *Smn^2B/-^* mice will serve as one of the very few models of microvesicular steatohepatitis for both adult and pediatric population. Like other models, the *Smn^2B/-^* mice are not representative of the complete NAFLD/NASH spectrum and features. Nevertheless, *Smn^2B/-^* mice offer a reliable, low-cost model to identify molecular players in pathogenesis of NAFLD.

## Results

### NAFLD in Smn^2B/-^ mice is prevented by gene therapy

We have previously identified microvesicular steatosis and dyslipidemia in *Smn^2B/-^* mice, occurring in the span of a few days, typically between postnatal day (P) 9 and P13[8]. The microvesicular steatosis and dyslipidemia in *Smn^2B/-^* mice is directly due to Smn depletion as it can be completely prevented by gene therapy using intravenous injection of the scAAV9-CB-SMN vector to induce exogenous expression (Fig 1A-F). Of note, the dose of vector (5 x 10^10^ VG) used for gene therapy leads to vector biodistribution mainly in peripheral tissues including liver, spleen and skeletal musculature[14]. Next, we sought to investigate the severity, functional consequences and mechanisms underpinning the NAFLD in these mice. Plasma levels of serum transaminases alanine aminotransferase (ALT) and aspartate aminotransferase (AST), markers of liver damage, were mildly elevated in P19 *Smn^2B/-^* mice (Fig 1G,H). However, plasma alkaline phosphatase (ALP) remained normal, but hepatic ALP staining was enhanced in livers from symptomatic *Smn^2B/-^* mice (Fig 1I-M). Muscular dystrophy patients can exhibit elevated serum transaminase levels, making skeletal muscle a potential source[15]. However, muscles from *Smn^2B/-^* mice are not degenerating[16], eliminating the possibility that they could be a source of transaminase. Altogether, our data is therefore indicative of liver damage. An active apoptotic process occurs as indicated by increased transcript levels for multiple cell death genes such as Fas receptor (*FasR*), TNF receptor superfamily member 1A (*TNFR1*), BCL2 associated X protein (*Bax*), and tumor protein p53 (*p53*) (Fig 1N), together with increased caspase 3 staining in livers of P17-19 *Smn^2B/-^* mice (Fig 1P,Q). The hepatic apoptosis appears to be P53-dependent, as expression of classical targets of P53[17], p21 and *Mdm2*, were strongly upregulated (Fig 1O). Finally, P17-19 *Smn^2B/-^* mice did not display any signs of liver fibrosis (Fig 1R-U). This analysis shows that *Smn^2B/-^* mice have steatohepatitis and hepatic cell death, but in the absence of significant fibrosis.

**Fig. 1.**
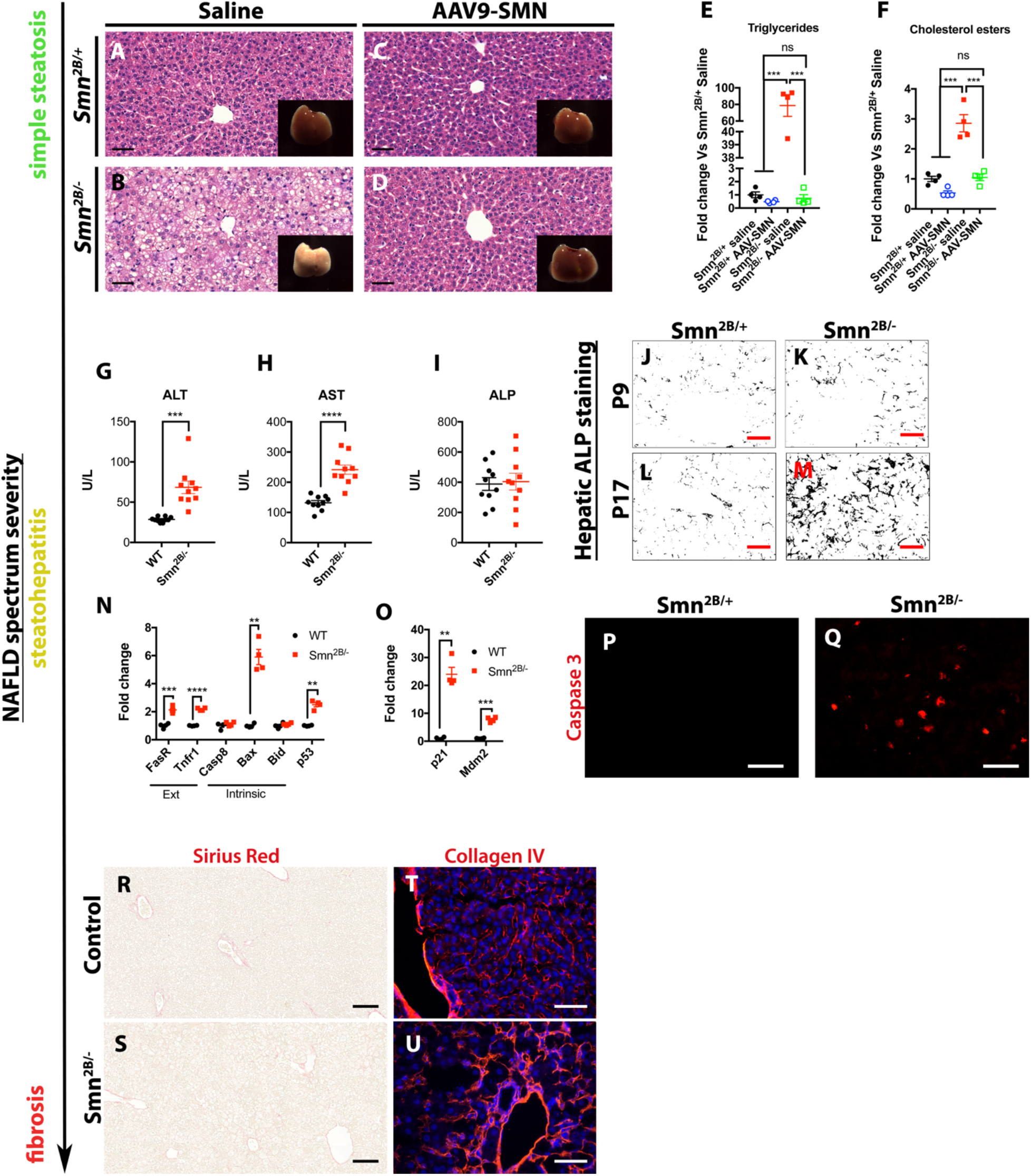
Symptomatic *Smn^2B/-^* mice suffer from significant liver damage without fibrosis. (A-F) Microvesicular steatosis is evident (40X, H&E staining) at P19 effectively prevented by systemic AAV9-SMN injection at P1. (G-I) Elevation ALT and AST, but not ALP in the plasma of *Smn^2B/-^* mice at P19. (J-M) Enhanced ALP staining (200X) in P17, but not P9, *Smn^2B/-^* livers. (N) FasR, TNFR1, Bax, p53, as well as p53 transcriptional targets (O) p21 and Mdm2 transcripts were significantly increased. (P-Q) Increased caspase 3 punctae in P17 *Smn^2B/-^* livers (100X). (R-U) Representative 20X and 200X images of Sirius red and collagen IV staining show no significant hepatic fibrosis in P17-19 *Smn^2B/-^* mice. QPCR data were normalized with SDHA and PolJ (H-I). Scale bar represents (A-D) 50 μm and (J-M, P-Q, R-U) 100 μm. N value for each experiment is as follows: N = 10 for G-I, 5 for J-M, 4 for A-F and N-O, 3 for P-Q, 5 for R-S, 3 for T-U. Statistical analysis were one-way ANOVA with Tukey’s multiple comparison test for E-F and unpaired twosided student’s t-test for G-I and N-O, *P* ≤ 0.05 for *, *P* ≤ 0.01 for **, *P* ≤ 0.001 for *** and *P* ≤ 0.0001 for ****)

### NAFLD in Smn^2B/-^ mice leads to alterations in multiple physiological processes

We next sought to identify whether liver damage in SMA translated into functional sequelae using important and translatable clinical readouts. Total protein and albumin were reduced in the plasma of P19 *Smn^2B/-^* mice (Fig 2A,B). Immune dysregulation has previously been identified in SMA model mice[18–20], and we identified a significant reduction in expression of many complement genes (Fig 2C). Abnormal blood clots have been previously reported in SMA[21, 22], and here we found altered transcript levels of genes involved in hemostasis (Fig 2D).

**Fig. 2.**
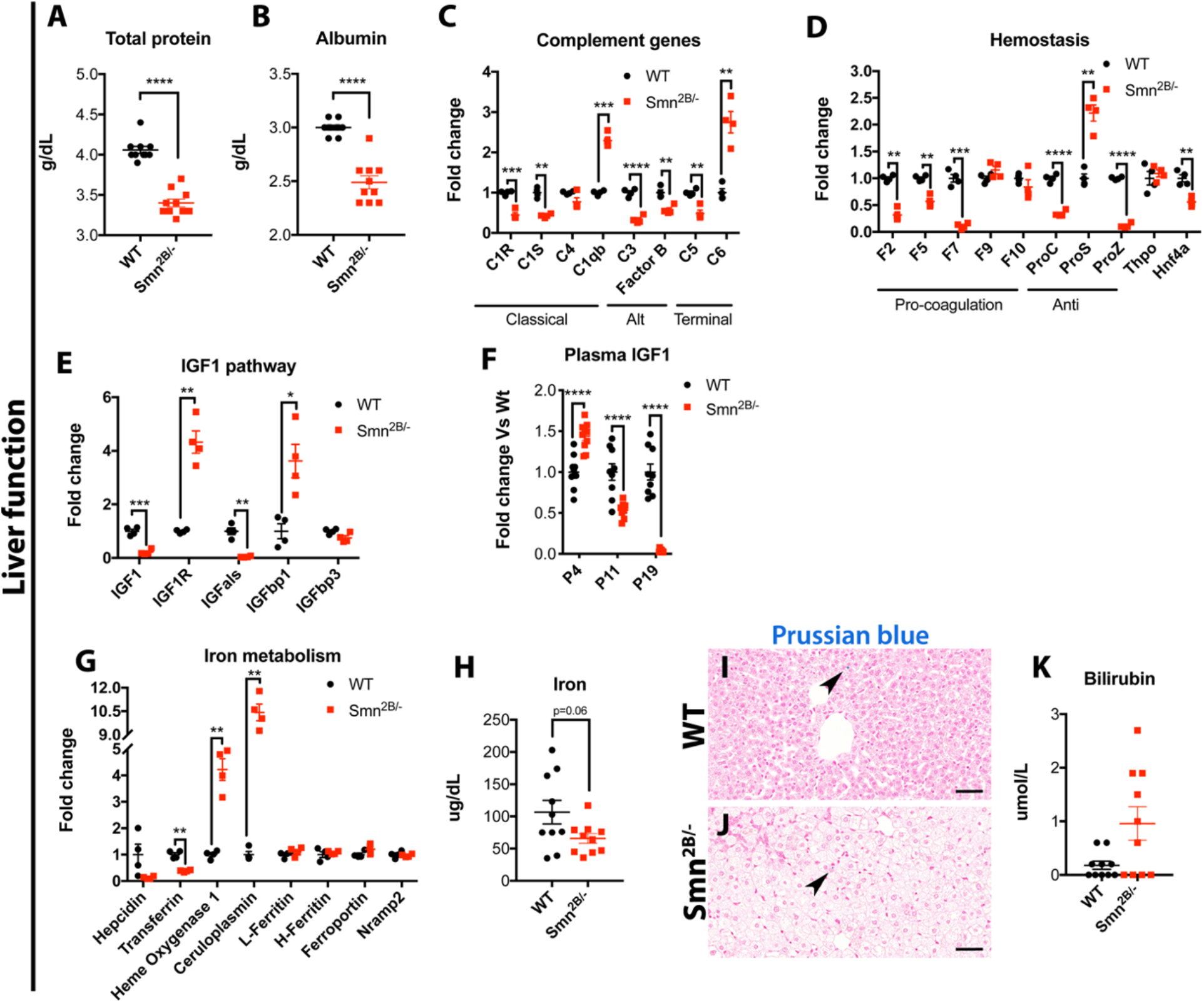
Liver functional deficits in multiple pathways in symptomatic *Smn^2B/-^* mice. (A,B) Low levels of total protein and albumin in plasma from P19 *Smn^2B/-^* mice. (C-E) Major alterations in levels of transcripts for complement, hemostasis and IGF1 pathway components in livers from P19 *Smn^2B/-^* mice. (F) Progressive depletion of IGF-1 hormone in the plasma from *Smn^2B/-^* mice. Iron metabolism genes are misregulated (G), and iron levels are reduced in plasma (H) but unchanged in liver (Prussian blue staining, 40X) (I,J). (K) A trend towards higher total bilirubin protein in the plasma in *Smn^2B/-^* mice. QPCR data were normalized with SDHA and PolJ (C,D, E). Scale bar represent 50 μm (I,J). (N value for each experiment is as follows: N = 8-10 for A-B, F, H, and K, 4 for C-E and G, 5 for I-J, unpaired two-sided student’s t-test for all except for (F) Two-way ANOVA with Sidak’s multiple comparison test, *P* ≤ 0.05 for *, *P* ≤ 0.01 for **, *P* ≤ 0.001 for *** and *P* ≤ 0.0001 for ****)

Liver is also a key source of growth factors, including insulin growth factor 1 (IGF1). We identified an important reduction in *Igf1* and insulin like growth factor binding protein acid labile subunit (*igfals*) transcript levels, and an upregulation of insulin like growth factor 1 receptor (*IGF1R*) and insulin like growth factor binding protein 1 (*IGFbp1*) transcript levels (Fig 2E). A remarkable and progressive reduction of plasma IGF1 protein was observed over time (Fig 2F). These data are consistent with previous reports in SMA[23–25].

We also identified many dysregulated transcripts for genes involved in iron metabolism, including *hepcidin*, a gene producing hepcidin protein that acts as a master regulator of iron levels[26], as well as *transferrin, heme oxygenase 1* and *ceruloplasmin* (Fig 2G). Plasma iron levels trended lower (Fig 2H) but hepatic stores appeared unaffected in *Smn^2B/-^* animals (Fig 2I,J). Finally, we have observed a trend towards higher levels of total bilirubin in the plasma, suggesting reduced efficacy of the hepatocytes to process it (Fig 2K). As such, it is likely that iron dysregulation in our model results from slow heme processing. Previous work has shown iron metabolism is affected by Smn depletion[22, 27]. Together, these findings demonstrate impairments in many functions of the liver.

### Identification of molecular mechanisms underpinning NAFLD in Smn^2B/-^ mice

To identify alterations in specific molecular pathways that could render SMA liver more susceptible to NAFLD, we undertook Tandem Mass Tagging (TMT) proteomic analysis of livers from pre-symptomatic P0 and P2 *Smn^2B/-^* mice compared to wild type, specifically to look for molecular changes present well before any overt pathology. We compartmentalized the data into biologically relevant subgroups based on the timing of altered protein abundance detection. This produced four subgroups, A, B, C and NS, where proteins in subgroup A (14% of the total IDs) represent those whose expression is already significantly altered at P0 but revert to wild type basal levels at P2. Subgroup NS (not altered at either P0 or P2) contained 65% of IDs (Fig 3A). We concluded that the proteins in these subgroups (A and NS) were therefore unlikely to be important for the NAFLD phenotype in the *Smn^2B/-^* mice. Conversely, proteins in subgroup B (unchanged at P0, but significantly changed at P2) included 11% of total proteins, while subgroup C (altered at both P0 and P2) including 10% of total proteins were of more interest. Analysis of subgroup B using BioLayout *Express*^3D^ and DAVID identified the mitochondrion cluster (increased protein expression) and the lipid metabolism cluster (decreased protein expression) (Fig 3B). A similar analysis of subgroup C identified clusters again associated with mitochondria (proteins significantly upregulated at both P0 and P2), extracellular signaling (proteins significantly decreased at both P0 and P2), and extracellular matrix proteins (significantly decreased at P0, however significantly increased at P2) (Fig 3C). To further refine potential pathways involved, we used ingenuity pathway analysis (IPA) software on proteins within subgroups B (Supp Fig 1) and C (Fig 3D). Of interest, the results from subgroup C revealed alterations in pathways related to “oxidative phosphorylation” (*p* = 6.35×10^−3^) and “mitochondrial dysfunction” (*p* = 1.11×10^−2^) (Fig 3D). Furthermore, IPA analysis identified “metabolism” (*p* = 3.53×10^−12^) and “homeostasis of lipids” (*p* = 1.68×10^−9^) as some of the top functional subgroupings perturbed in *Smn^2B/-^* liver at P0 (Fig 3E). Thus, this proteomic screen points towards mitochondrial dysfunction, a critical player in fatty acid clearance through *β*-oxidation.

**Fig. 3.**
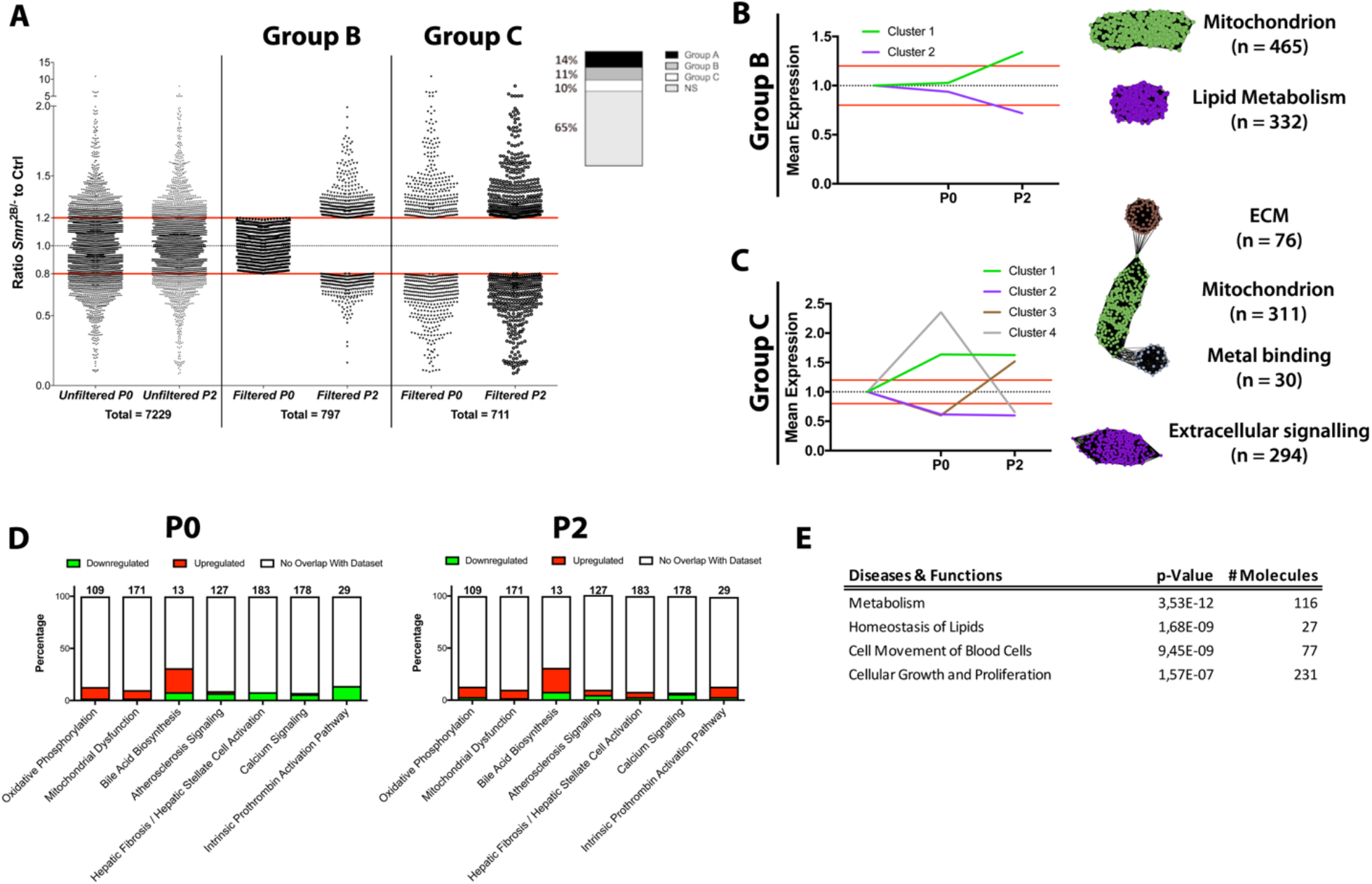
Proteomic analysis of P0 and P2 *Smn^2B/-^* livers identifies mitochondrial and lipid metabolism as prominent perturbations. (A) Scatterplots showing protein expression ratios of *Smn^2B/-^* to P0 wild type (control) liver. A 20% threshold altered expression was applied. Left column of paired scatterplots shows *Smn^2B/-^* to wild type ratios for 7229 proteins at birth (P0) and P2. Group B identified by filtering for proteins altered only at P2 in *Smn^2B/-^* livers (P0 = ns (0.8 ≤ x ≤ 1.2) and P2 = *p* ≤ 0.05 (x < 0.8 or x > 1.2)). Group C filters for proteins altered at P0 and at P2 in *Smn^2B/-^* (P0 and P2 = *p* ≤ 0.05 (x < 0.8 or x > 1.2). (B-C) Group B and Group C graphical representation of *Smn^2B/-^* to wild type ratio proteins at P0 and P2, left graph prior to clustering, right graphic post application of the MCL clustering algorithm (inflation value 2.2) analyzing coordinately expressed proteins. These are represented as mean ratio-change per cluster. In cluster visualization the proteins are spheres with correlation between them of r ≥ 0.9 indicated by black lines. Each identified cluster has a functional annotation with n number stating how many proteins are present within the cluster. (D) IPA top canonical pathways highlighting the main disrupted cascades in Group C data set at P0 (left) P2 (right). Stacked bar chart displays the percentage of proteins that were upregulated (red), downregulated (green), and proteins that did not overlap with our data set (white) in each canonical pathway. The numerical value at the top of each bar represents the total number of proteins in the canonical pathway. (E) Top diseases and functions linked to our Group C data set identified by IPA functional analysis. (see methods for comprehensive description of analysis)

### Assessment of mitochondrial function in livers of Smn^2B/-^ mice

Given the proteomic data findings and the possibility that impaired mitochondrial function could be driving NAFLD/NASH in *Smn^2B/-^* mice, we focused on mitochondrial content, structure and function. Oxidative phosphorylation complex protein levels are largely unchanged at P9 in liver tissue homogenate. However, the protein expression of SDHB (complex II), MTCO1 (complex IV) and ATP5A (complex V) were reduced in tissue homogenate of P19 *Smn^2B/-^* livers (Fig 4A,B), highlighting a potential depletion of mitochondrion number. A reduced mitochondrial density was confirmed by a lower activity of the citrate synthase enzyme at P19-21 (Fig 4C)[28]. Cursory ultrastructural analysis of mitochondria revealed no obvious gross alterations (Fig 4D-G). Surprisingly, high-resolution respirometry of isolated liver mitochondria from P19-21 *Smn^2B/-^* mice identified increased leak and ADP phosphorylation capacities when fueled by pyruvate, malate, and succinate (Fig 4H-L), or palmitoyl carnitine (Supp Fig 2). Interestingly, *Smn^2B/-^* mitochondrial function was similar to control mice at P9, a time point where hepatic fat accumulation is not readily observed[8]. Hepatic mitochondria from P9 *Smn^2B/-^* mice also exhibited an increase in reactive oxygen species (ROS) production (Fig 4M-Q). It is possible that the increased capacity for respiration in isolated mitochondria from P19 *Smn^2B/-^* mice is a compensatory mechanism to restore metabolic homeostasis and/or in response to low mitochondrial density. In addition, the enhanced ROS production could be responsible in part for hepatocyte damage and death (Fig 1G-Q). The increased capacity for fatty acid-supported respiration was consistent with the elevated levels of microsomal oxidation enzyme CYP4A (Fig 4R,S), known to be active upon *β*-oxidation overload[10, 29, 30]. Carnitine palmitoyltransferase I (CPT1), an enzyme responsible for shuttling long chain fatty acid into the mitochondria for *β-* oxidation, can be inhibited by malonyl-CoA, a product of *de novo* lipogenesis[30]. Such inhibition would lead to further fatty acid overspill in the microsomal oxidation pathway. We found CPT1 to have reduced activity in comparison to both wild type and *Smn^2B/+^* mice at P19 (Fig 4T). Overall, our results show that mitochondrial function of isolated mitochondria is normal or increased in the *Smn^2B/-^* mice, when oxidative processes are supported directly by substrates for complexes I and II. Given that CPT1 activity was decreased, it is possible that there is impaired formation of acyl carnitine species or inhibition of CPT1 activity, activation of proton leak and reduced uptake of long chain fatty acids for mitochondrial oxidation *in vivo*, further exacerbating hepatic steatosis.

**Fig. 4.**
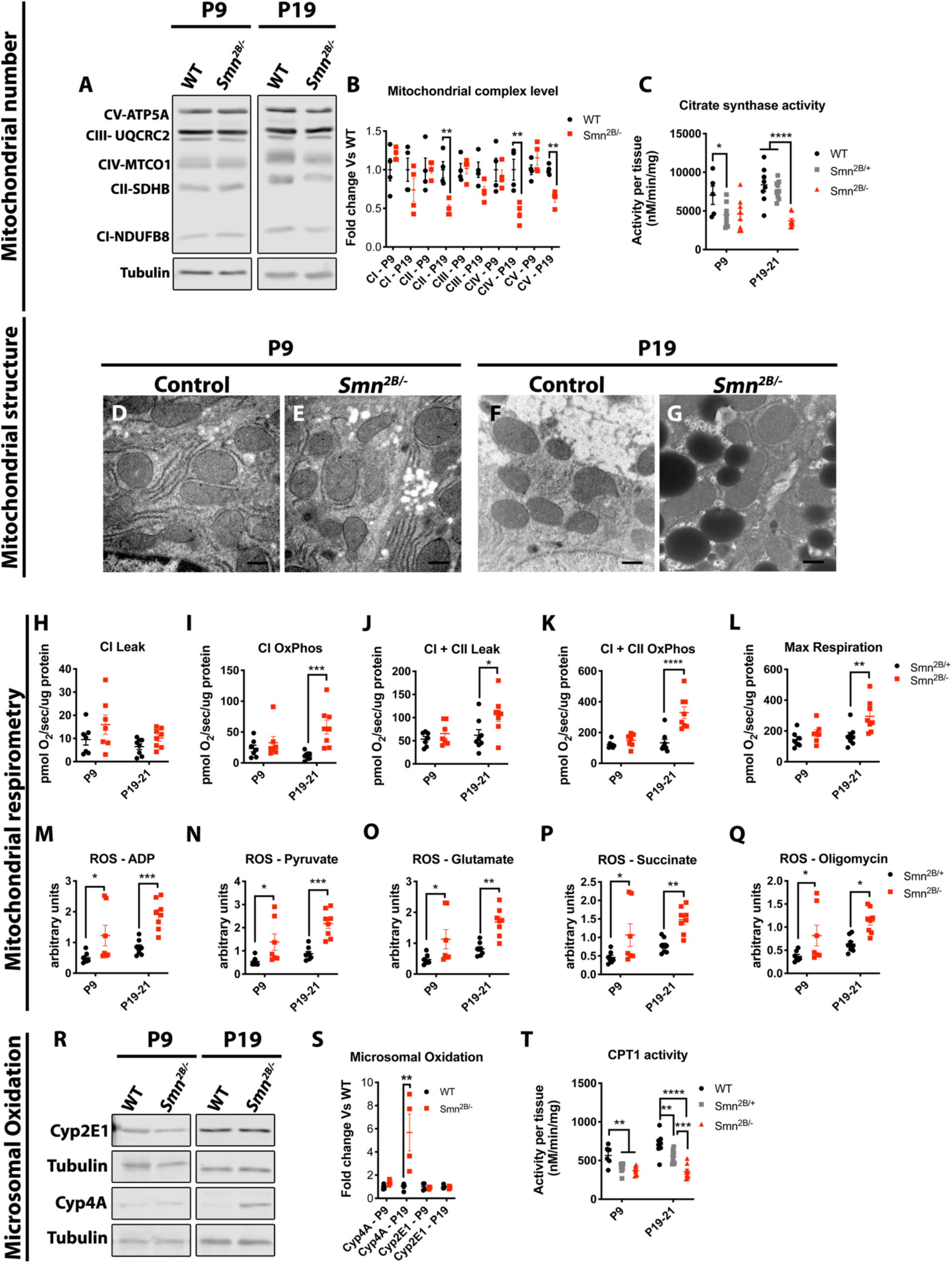
*Smn^2B/-^* liver mitochondria show increased β-oxidation and ROS production. (A,B) Western blot analysis of subunits of the mitochondrial complexes shows no significant change prior to hepatic fat accumulation at P9 but shows significant down-regulation of CII, CIV and CV subunit proteins in P19 *Smn^2B/-^* liver homogenates. (C) Lower citrate synthase activity in livers of P19-21 *Smn^2B/-^* mice, suggests decreased mitochondrial density per mg of tissue. (D-G) Mitochondrial structure appears relatively spared at both P9 and P19 in *Smn^2B/-^* livers. (H-L) High resolution respirometry of *Smn^2B/-^* hepatic mitochondria shows increased leak and higher respiratory capacity at P19 but not at P9 in comparison to *Smn^2B/+^* hepatic mitochondria. (M-Q) *Smn^2B/-^* hepatic mitochondria had an increase in ROS production during most respiratory states in comparison to *Smn^2B/+^* mitochondria. (R,S) Increased expression of CYP4A is evident in P19 *Smn^2B/-^* livers. (T) Reduced CPT1 activity is present in livers of *Smn^2B/-^* mice compared to WT. (N value for each experiment is as follows: N = 3-4 for A-B, & R-S, 1 for D-G, 6-9 for C, H-Q & T, Two-way ANOVA with Sidak’s multiple comparison for all except C & T two-way ANOVA with Tukey’s multiple comparison tests, *P* ≤ 0.05 for *, *P* ≤ 0.01 for **, *P* ≤ 0.001 for *** and *P* ≤ 0.0001 for ****)

### Hormonal contribution to NAFLD in Smn^2B/-^ mice

Insulin insensitivity plays a major role in the development of NAFLD. *Smn^2B/-^* mice show abnormal glucose handling in intra-peritoneal glucose tolerance test[31] and small cohort of SMA patients showed susceptibility to insulin resistance[32]. Surprisingly, the *Smn^2B/-^* mice show sustained hypoglycemia with age in a normoinsulinemic state and a trend towards diminished C-peptide production at P19 (Fig 5A-C). This is in line with relatively low HbA1C observed in SMA patients[8]. Due to their small size and age, hyperinsulinemic clamp is not feasible to further assess insulin sensitivity. Alternatively, we also noted a progressive elevation of plasma glucagon levels, which was first evident at P11 in *Smn^2B/-^* mice (Fig 5D). This increase in glucagon likely results from the increase in alpha-cell number in *Smn^2B/-^* pancreas[31] and/or low glucose. Glucagon signaling mediates some of its effects through the phosphorylation of Creb, which leads to expression of the gluconeogenic program[33]. We observed increased phospho-Creb levels in livers of P19 *Smn^2B/-^* mice (Fig 5E). Interestingly, there is a robust increase in the levels of GLP-1 (Fig 5F), another byproduct of proglucagon, produced in the gastrointestinal tract. Enhanced glucagon levels/signaling lead to glycogenolysis and gluconeogenesis in the liver, and lipolysis in the white adipose tissue to increase energetic substrate availability in the bloodstream[34]. Pathological glucagon signaling could lead to energy substrate overload in the blood, and subsequent stimulation of the liver to restore homeostasis via uptake of these substrates, including lipids. While limited change was identified in the time frame of acute fat accumulation in the liver on pathology (between P11-13[8]) (Fig 5I-L), we observed eventual hepatic glycogen depletion (Fig 5G-H), a trend towards adipocyte size reduction (Fig 5M-S) and increased NEFA (Fig 5T), a direct product of lipolysis, in the blood. These findings are consistent with enhanced glucagon signaling. More particularly, NEFA level was readily observable at P11 and worsened over time in comparison to control (Fig 5T). Triglyceride levels followed a similar progression, albeit in a delayed fashion (Fig 5U). Altogether, these findings point to a fatty substrate overload in the blood as a consequence of glucagon pathway activation.

**Fig. 5.**
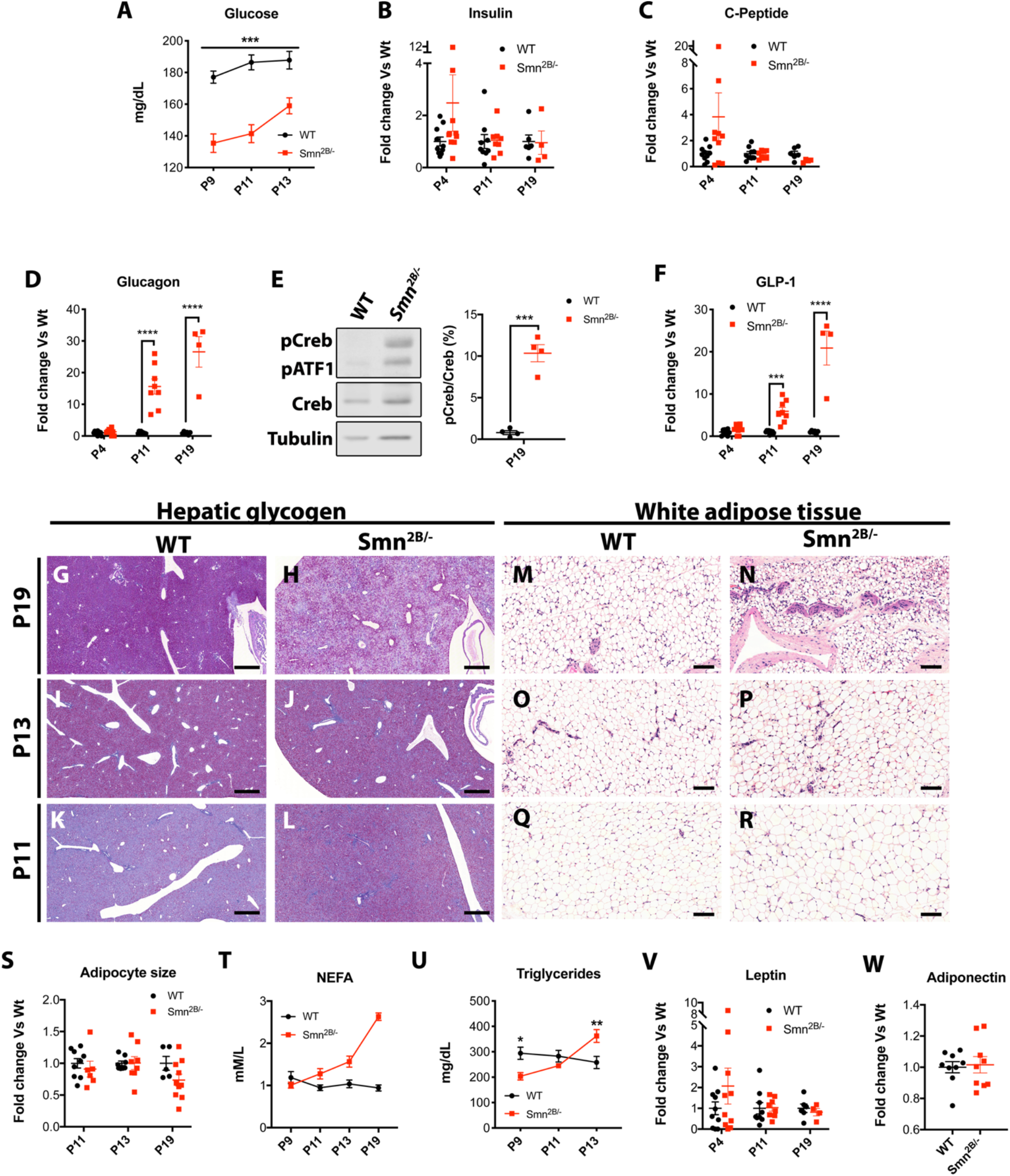
Hyperglucagonemia leads to increased substrate release in the plasma of *Smn^2B/-^* mice. (A) Plasma glucose was lower throughout P9 to P13 in *Smn^2B/-^* mice in comparison to wild type mice. (B) Insulin levels was relatively maintained throughout the *Smn^2B/-^* mice lifespan. (C) Trend towards diminished C-Peptide production is only seen at P19 in *Smn^2B/-^* mice. (D) Progressive elevation of plasma glucagon occurs in *Smn^2B/-^* mice with a ~15-fold increase by P11. (E) Western blot analysis shows a 10-fold increase in phospho-Creb, a downstream molecular event of glucagon activation, in P19 *Smn^2B/-^* livers in comparison to WT. (F) Plasma GLP-1, a product of the cleavage of proglucagon, is altered in a similar fashion. (G-L) PAS stained liver sections (5X) at P19, P13 and P11 reveal glycogen depletion in P19 *Smn^2B/-^* mice (G,H), but not at P13 (I,J) and P11 (K,L). (M-R) H&E sections (20X) of subcutaneous adipose tissue show a trend toward reduction in adipocyte size at P19 *Smn^2B/-^* mice (M,N,S) but not at P13 (O,P,S) and P11 (Q-S). (T) Plasma NEFA progressively increase from P9 to P19, concordant with increased lipolysis of white adipose tissue. (U) Plasma triglyceride quantification showed a similar trend as NEFA, albeit in a delayed fashion. (V-W) Adipokines leptin and adiponectin levels remained relatively unchanged. Scale bar represents 500 μm in G-L and 100 μm in M-R. (N value for each experiment is as follows: N = 8-10 in A, T-U, W, 8-10 for P4, P11 and 4-6 at P19 in (B-D, F, V), 4 in E, 5 for G-L, 5-10 for M-S, Two-way ANOVA with Sidak’s multiple comparisons test for (A-D, F, S, U, V), unpaired two-sided student’s t-test for E and W, note that no statistical analysis were performed on (T) given results obtained through different techniques for P19, *P* ≤ 0.001 for *** and *P* ≤ 0.0001 for ****)

We further investigated adipocytic hormones (i.e. leptin, adiponectin), which are known to play a role in NAFLD/NASH progression and fibrosis[10]. We did not observe any changes in leptin or adiponectin in *Smn^2B/-^* mice (Fig 5V-W). Other hormones from the gastrointestinal tract (ghrelin, GIP, PYY), pancreas (PP, amylin) and adipocyte (resistin) hormones did not show a consistent pattern of misregulation (Supp Fig 3).

## Discussion

We systematically characterized typical features of NAFLD development to better understand the metabolic abnormalities in *Smn^2B/-^* mice. These mice display many positive features of NALFD. They develop microvesicular steatohepatitis without fibrosis within two weeks of life, with increased serum markers of liver damage, and hepatocyte cell death. They also display significant dyslipidemia[8], peripheral lipolysis, functional hepatic deficits, alterations in mitochondrial function, evidence of involvement of alternative oxidative pathways and ROS production in the liver. All of these features have been observed in NAFLD[10]. Nevertheless, the *Smn^2B/-^* mouse as a model for NAFLD also has some limitations. They do not develop fibrosis, a component that is seen in NASH patients[10]. *Smn^2B/-^* mice also lose weight due to their neurological condition, display low blood sugar and normal insulin levels. In contrast, most NAFLD patients have metabolic syndrome, which includes features of obesity, dyslipidemia, insulin resistance, hyperglycemia, hyperinsulinemia[10]. Note that it is unclear whether the *Smn^2B/-^* mice develop insulin resistance as gold standard studies are not logistically feasible in this model, but SMA patients appear prone to insulin resistance[32]. *Smn^2B/-^* mice show some difficulties in glucose handling during intra-peritoneal glucose tolerance challenge[31]. In comparison, some popular NAFLD models also show limitations. For example, the methionine and choline deficient diet (MCD) model does not exhibit any of the metabolic features[10–12, 35]. The *ob/ob* and *db/db* mutant mice, which display altered leptin signaling, have metabolic features but no inflammation or fibrosis[10–12]. A second challenge, such as high fat diet (HFD) or MCD, is needed to develop these features in the *db/db* mouse[36]. The HFD diet appears to result in all features of the NAFLD spectrum, however, fibrosis is minimal and can take up to 36-50 weeks to develop[37]. Although not perfect either, the *Smn^2B/-^* mice could provide an efficient mouse model of NAFLD with microvesicular steatosis, short turnaround time (about 14 days[8]), and limited need for special diet, which would speed up identification of molecular targets. The fast steatosis phenotype, paired with the fact that no special and expensive diet is required, make it a cost-effective option. In fact, introduction of high fat diet did not drastically worsen the overall metabolic phenotype of the *Smn^2B/-^* mice[38]. In addition, our study made use of both male and female mice, unlike other mouse models where males are predominantly used[39]. Needless to say, the *Smn^2B/-^* mice would allow for a different outlook on molecular players and organ system involvement in comparison to current available models of NAFLD. Indeed, the *Smn^2B/-^* mice could act as one of the few mouse models for pediatric NAFLD and/or microvesicular steatosis[39, 40]. To our knowledge, all current NAFLD models mostly display macrovesicular steatosis, apart from the *Acox^-/-^* mice, which develop predominantly microvesicular steatosis[41].

Microvesicular steatosis is present in all *Smn^2B/-^* mice. In the pediatric population, microvesicular steatosis is generally clinically associated with inherited metabolic disorders and fatty acid oxidation defects[42]. In contrast, it is only present in a minority (10%) of adult NAFLD patients and associated with more severe disease[43]. Strikingly, diffuse microvesicular steatosis, as seen in our model, can present with encephalopathy and liver failure due to underlying severe mitochondrial *β*-oxidation dysfunction[43, 44]. Interestingly, one SMA patient had to undergo liver transplant due to acute liver failure post-operatively and had both macro and microvesicular steatosis[6]. More recently, we found that of 8 SMA liver biopsies, 3 (37.5%) had microvesicular steatosis, in comparison to the expected 0.7%[45] of patients with liver steatosis in 2-4 year old range in the normal population[8]. Additionally, a fatty acid oxidation phenotype was previously described by the presence of dicarboxylic aciduria in some SMA patients[46], which may be caused by activation of microsomal oxidation as shown in the present study. We found no evidence of a β-oxidation deficit in our model using high resolution respirometry in isolated liver mitochondria. On the contrary, it appears that the isolated mitochondria have enhanced capacity, perhaps reflective of a compensatory reaction. We suggest that the compensation is due to both the reduced mitochondrial density as well as the increase triglyceride storage in the liver of *Smn^2B/-^* mice. To note, CPT-1 activity was much reduced, which may reduce transport of long-chain fatty acid for oxidation. A decrease in CPT1 activity can in turn lead to increased fatty infiltration and liver damage. Nevertheless, our proteomic screen identified alterations in two important clusters, namely mitochondria and lipid metabolism, close to birth, and well before any overt neurological or hepatic pathology develops. Interestingly, “mitochondrial pathway components” are often represented in “omic” data of SMN depleted tissue, including motor neurons[47], diaphragmatic NMJs[48], hippocampal synapses[49], isolated motor neurons[50], induced pluripotent stem cell motor neurons[51], or directly related to motor neuron vulnerability[52]. Abnormal mitochondrial findings have previously been reported in cell culture, SMA models, and SMA patients[52–55]. Additionally, it is also part of NAFLD/NASH pathogenesis[10]. As such, additional investigation will be required to refine mitochondrial defects in this model and how it can relate to NAFLD/NASH.

From our analysis, we conclude that NAFLD development in *Smn^2B/-^* mice is multifactorial. The proposed mechanism underpinning the defects is illustrated in Fig 6. We propose that the initial event leading to fatty acid dysregulation in the liver likely stems from abnormal glucose homeostasis. Hyperglucagonemia is induced early in *Smn^2B/-^* mice in response to low blood glucose in the bloodstream or from the pathological overpopulation of alpha cells in the pancreas[31]. Surprisingly, glucose levels in the *Smn^2B/-^* mice are reduced as early as P9. The glucose level remains low but is sustained, likely due to gluconeogenesis. Eventually, gluconeogenesis fails due to depleted glycogen storage in P19 *Smn^2B/-^* mice, leading to a sudden drop in glucose level in the blood[8]. Simultaneously, lipolysis of white adipose tissue, a by-product of glucagon signaling, is induced to ensure availability of energy substrate, represented by a progressive increase in NEFA from P9 to P19 in *Smn^2B/-^* mice. This leads to increased fatty substrates in the bloodstream, which precede or coincide with muscle denervation. Skeletal muscle, a major consumer of energy substrates when innervated and fully functional, will have a diminished requirement for energy as denervation renders it non-functional in SMA. As such, this leads to overload of fatty energy substrates in the circulation. Eventually, the susceptible SMA liver will take up the lipid substrates for storage in an attempt to restore homeostasis, which in turn leads to liver steatosis. Pathological fat storage could spill over to the muscle compartment once the liver has reached saturation, which is consistent with our previous description of lipid droplets on ultrastructural analysis of skeletal muscle of *Smn^2B/-^* mice[56]. Finally, enhanced ROS production from mitochondrial oxidation leads to hepatic damage and functional deficits.

**Fig. 6.**
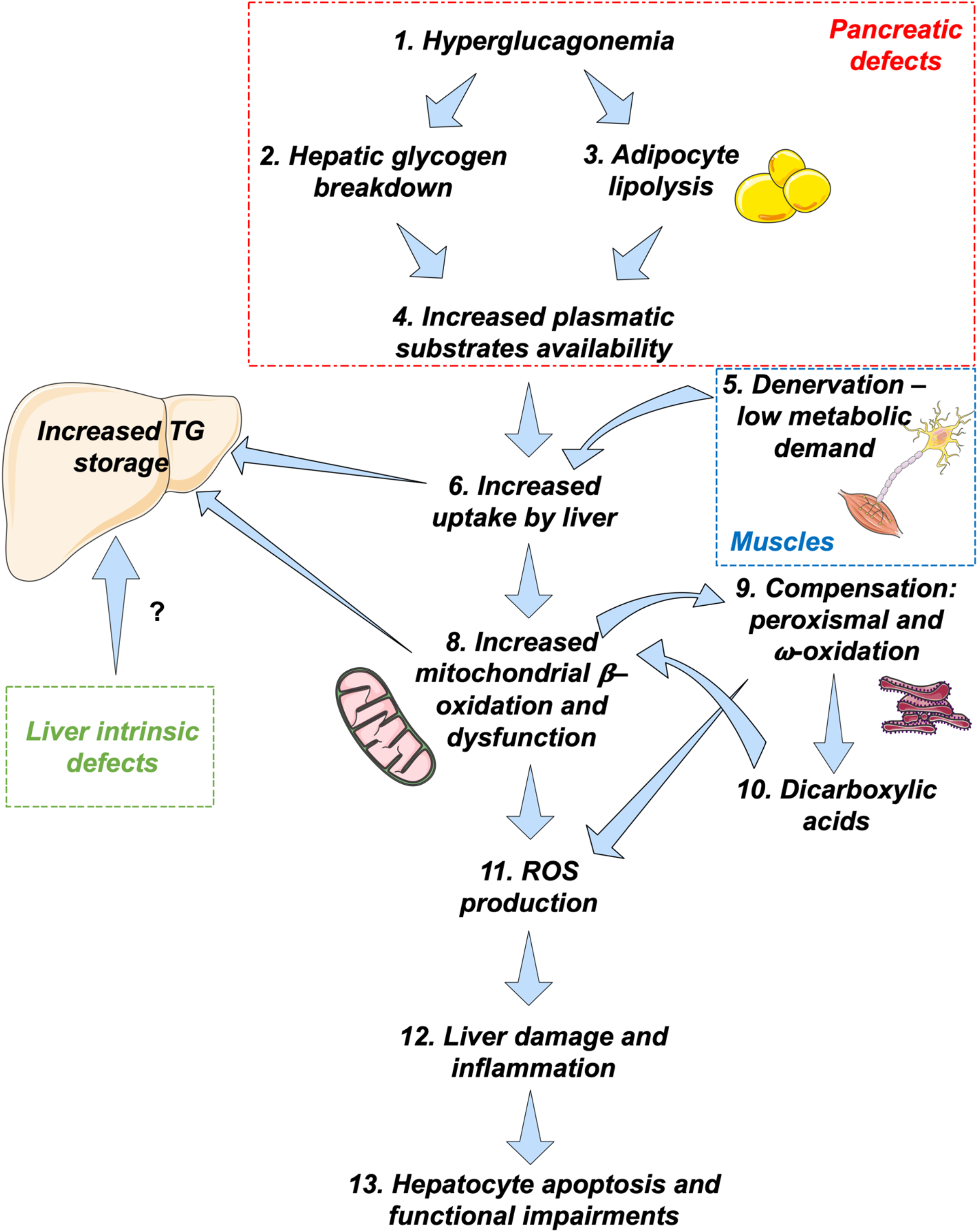
Schematic summarizing the findings of the present study. Undefined glucose abnormalities lead to hyperglucagonemia leading to hepatic glycogen breakdown and adipocyte lipolysis. This results in increased plasma energy substrate availability prior or concomitantly to muscle denervation, the major user of energy in the blood. This leads to overload of fatty substrates in the blood, which the liver takes up to restore homeostasis, leading to steatosis. To compensate and dispose the unnecessary lipids, mitochondrial oxidation is increased to burn excess lipids, which eventually becomes overloaded and requires alternative peroxisomal and microsomal oxidation pathway. Such a compensation leads to increased ROS production, liver damage, hepatocyte apoptosis and eventually functional impairment. The schematic art pieces used in this figure were provided by Servier Medical art https://smart.servier.com. Servier Medical Art by Servier is licensed under a Creative Commons Attribution 3.0 Unported License

Altogether, *Smn^2B/-^* mice will provide an appropriate NAFLD model, more particularly for pediatric and microvesicular pathogenesis. It can be leveraged for high throughput identification of molecular pathways involved in NAFLD due to the fast onset of the phenotype (less than 2 weeks) and the lack of a required diet, making it a cost-effective option in the study of NAFLD pathogenesis.

## Materials and Methods

### Study design

We have recently identified the increased prevalence dyslipidemia and fatty liver[8] in SMA patients and SMA mouse models. This sparked a project with the following 2 pre-specified objectives: (1) Identify consequences of the fatty acid defect, (2) Identify the etiology of these defects and how it relates to NAFLD pathogenesis. As it pertains to this manuscript, the etiologies of the defects were suspected on prior experience with this SMA mouse model and NAFLD literature. It included denervation[8], liver-intrinsic defects, mitochondrion, and external factors (other organs). Serum analysis and lipid quantification were outsourced, and, thus, analyses were performed in a blinded fashion. N number are described in each figure legend. Statistical approach is as described below and in figure captions. Collaboration between laboratories of Kothary and Parson and colleagues occurred mid-project, given overlapping results that were converging. Hence, the resulting manuscript offers data that have been concordant in two independent laboratories (albeit using different experimental paradigms).

### Mouse models and treatments

The *Smn^2B/-^* (wild type C57BL/6J background)[9] mouse lines were housed at the University of Ottawa Animal Facility and cared for according to the Canadian Council on Animal Care. Experimentation and breeding were performed under protocol OHRI-1948 and OHRI-1927. *Smn^+/-^* mice were crossed to *Smn^2B/2B^* mice to obtain *Smn^2B/+^* and *Smn^2B/-^* animals. C57BL/6J wild type mice were bred separately. All experiments using mice in the UK were performed in accordance with the licensing procedures authorized by the UK Home Office (Animal Scientific Procedures Act 1986). All tissues in the Kothary laboratory were collected while mice were fed *ad libitum*. Tissues undergoing biochemical analysis in the Kothary laboratory were collected between 9 and 11 AM to limit the effect of the circadian rhythm.

### scAAV9-CB-SMN production and injection

scAAV9-CB-SMN vectors were produced at the Bertarelli Platform for Gene Therapy in EPFL (Switzerland), using a construct similar to the one described in [57]. The self-complementary scAAV9-CB-SMN vector was produced by calcium phosphate transfection of HEK293-AAV cells (Agilent) with pAAV-CB-SMN[57] and pDF9 plasmids. Briefly, the vector was purified from the cell lysate using a iodixanol density gradient followed by anion exchange chromatography (HiTrap Q-FF column, GE Healthcare). The scAAV9-CB-SMN vector was finally resuspended and concentrated in DPBS on a centrifugal filter unit (Amicon^®^ Ultra-15, Millipore). The titer of the vector suspensions was determined by qPCR using an amplicon located in the inverted terminal repeats as described in[58]. The obtained titers of the scAAV9-CB-SMN vectors were 9.6^13^ VG/mL and 3.0^13^ VG/mL. *Smn^2B/-^* and *Smn^2B/+^* mice were injected with 5 x 10^10^ VG of the AAV9-CB-SMN viral vector at P1 through the facial vein and the mice were then allowed to age until P19.

### Gross morphology, tissue processing and staining of animal tissues

Livers and white adipose tissue were fixed in formalin (1:10 dilution buffered, from Protocol, cat #245-684) for 24-48 h or 72 h (white adipose tissue) at 4°C and then transferred in 70% ethanol at 4°C until processing. All samples used for histological assessment were processed at the University of Ottawa (Department of Pathology and Laboratory Medicine) and embedded in wax using a LOGOS microwave hybrid tissue processor. Paraffin block tissues were cut with a microtome at 3-4 μm thickness. Hematoxylin & eosin (H&E) staining was performed using a Leica autostainer XL. Periodic acid-Schiff (PAS), Prussian blue, oil red O and Sirius red staining were performed using standard methods. Tissue undergoing immunofluorescence were processed in Dr. Parson’s laboratory. The tissues for immunofluorescence staining were sectioned (5 μm) on a cryostat (Leica, CM3050 S) or a microtome (Leica, RM2125 RTS). Liver sections were stained for caspase 3 (Abcam Ab13847 1:100) and collagen IV (Millipore AB756P 1:100). Antigen retrieval was performed to visualize Casp3. Briefly, air dried sections were quickly washed in PBS, then submerged into pre-warmed Antigen Retrieval Buffer and placed into water bath set at 90°C for 40 min (Caspase 3). The sections were removed from the bath but left submerged in the buffer allowing them to cool down but not dry out. After approximately 30 min, the slides were quickly washed in PBS and subjected to the traditional IHC staining protocol. Acquisition of signal was either obtained by slide scanning with a MIRAX MIDI digital slide scanner (Zeiss) and images acquired using 3DHISTECH Panoramic Viewer 1.15.4/CaseViewer 2.1 or directly captured using a Nikon eclipse e400 microscope (x10, x20 or x40 objective) and its images captured using QICAM Fast 1394 camera and Improvision Velocity 4 image capture software.

### Total ALP *in situ* assay

Liver sections on slides were washed in PBS twice for 5 min each, followed by three washes in NTMT (100 mM NaCl, 100 mM Tris-HCl pH 9.5, 50 mM MgCl_2_, and 1% Tween-20) for 10 min each. After incubation in color reaction solution (NTB, BCIP, and NTMT buffer), sections were washed in NTMT twice for 10 min each, followed by two washes in PBS for 10 min each. Sections were post-fixed in 4% PFA for 30 min, washed in PBS twice for 10 min each, and slides mounted in 90% glycerol.

### Gene expression studies

RNA from liver and spinal cord were extracted using Qiagen RNeasy Mini kit and reverse transcribed using RT^2^ first strand kit according to manufacturer’s protocol. A complete list of primers is available in the supplementary material (Supplementary Table 1). A standard curve was performed for each primer set to ensure their efficiencies. Each QPCR reaction contained equal amount of cDNA, Evagreen SyBR (Biorad), RNase/DNase-free water and appropriate primers (100-200 nM or according to PrimePCR protocol) in a final volume of 25 μl or 20 μl (for primePCR primers). To confirm amplicon specificity, a melting curve analysis was performed. Two negative controls were included in every QPCR plate and consisted of water in lieu of cDNA. QPCR results were quantified using 2^−ΔΔCt^ method. Results were normalized with 2 genes (mentioned in each figure legend containing QPCR data) identified as appropriate stable internal reference given M value below 0.5 and coefficient of variance below 0.25.

### Immunoblotting

Total protein lysate was collected by homogenization of flash frozen liver in RIPA lysis buffer (Cell Signaling). Protein concentrations were determined using the Bradford assay (Bio-Rad) or BCA assay. Protein extracts were subjected to sodium dodecyl sulfate polyacrylamide gel electrophoresis and examined by immunoblot, as previously described[59] with modified blocking conditions where Odyssey blocking buffer (Li-Cor 927-40000) replaced 5% milk. Primary antibodies used were as follows: pCreb (Ser133) (Cell Signaling 9198, 1:1000), Creb (Cell Signaling 9104, 1:1000), MitoOxphos (Abcam, ab110413 - 1:250), alpha-tubulin (Abcam, ab4074 - 1:2500-5000 and Calbiochem, CP06 1:10000), Cyp4A (Abcam, ab3573 - 1:1000), Cyp2E1 (Abcam, ab28146 - 1:2500). Secondary antibodies used were IRDye (Li-Cor) 680 or 800 (Li-Cor - 1:10000 to 1:20000). Signals were detected with Odyssey CLx (Li-Cor). Results were normalized to total protein or tubulin. Full western blots can be visualized in Supp Fig 4.

### Transmission Electron Microscopy

Electron microscopy was performed as previously described[56].

### High-resolution respirometry and mitochondrial enzymatic assays

Livers were excised from P9 and P19-21 *Smn^2B/+^* and *Smn^2B/-^* mice. Mitochondria were isolated using a slightly modified version from[60]. Briefly, livers were washed in IB_C_ buffer (see protocol[60]) and then minced and resuspended into 3 (P19) or 2 ml (P9) of IB_C_ buffer. Liver pieces were then transferred to a glass-Teflon homogenizer for homogenization using electric rotator. The homogenates were then centrifuged at 800g for 10 min at 4°C, supernatant was transferred to a new tube and centrifuged again 8600g 10 mins at 4°C, where pellet was resuspended in half initial volume of IB_C_ buffer. This process was repeated once. Mitochondria were indirectly quantified by Bradford assay. 700 ug (P19-21) and 500 ug (P9) of mitochondria were then introduced in the high-resolution respirometer (O2K; Oroboros, Austria) for respirometry measurements. The list and order of substrates and compounds introduced in the chamber for each protocol can be found in Supplementary Table 2 and 3. The substrates and compounds were added to the chamber after mitochondria reached steady state. Quantification was performed using the Oroboros software.

#### Citrate synthase and CPT1 activity

Enzyme activity for citrate synthase (CS) and CPT1 was determined as previously described with some modifications[61]. Briefly, tissue was weighed and homogenised in ice-cold homogenisation buffer (25 mM TRIS-HCL pH7.8, 1 mM EDTA, 2 mM MgCL_2_, 50 mM KCL, 0.50% Triton X-100) using modified Dounce homogenization with a pestle attached to a rotor. Homogenates were centrifuged at 14,000 x g for 10 min at 4°C, and the supernatant was collected. The assay was performed using the BioTek Synergy 96-well microplate reading spectrophotometer at room temperature. CS activity was determined by measuring absorbance at 412 nm in 50 mM Tris-HCl (pH 8.0) with 0.2 mM DTNB, 0.1 mM acetyl-coA and 0.25 mM oxaloacetate. Rate of absorbance change, and path length of each well was determined using BioGen 5.0. The enzyme activities were calculated using the extinction factor, 13.6 mM-1cm-1 for CS. For CPT1 enzymatic assay. CPT1 activity was determined by measuring absorbance at 412 nm in 50 mM Tris-HCl pH8.0 with 0.2 mM DTNB in a buffer containing 150 mM KCl, 0.1 mM palmitoyl-CoA and 0.25 mM l-carnitine. Enzymatic activity was reported as the activity per mg of tissue.

### Lipid quantification

Tissues were extracted and flash frozen. When required, tissues were pooled to obtain 100 mg. Tissue lipid analysis for quantification and profiles were performed at the Vanderbilt Mouse Metabolic Phenotyping Center. Briefly, lipids were extracted using the method of Folch-Lees[62]. The extracts were filtered, and lipids recovered in the chloroform phase. Individual lipid classes were separated by thin layer chromatography using Silica Gel 60 A plates developed in petroleum ether, ethyl ether, acetic acid (80:20:1) and visualized by rhodamine 6G. Phospholipids, diglycerides, triglycerides and cholesteryl esters were scraped from the plates and methylated using BF3/methanol as described in [63]. The methylated fatty acids were extracted and analyzed by gas chromatography. Gas chromatographic analyses were performed on an Agilent 7890A gas chromatograph equipped with flame ionization detectors, a capillary column (SP2380, 0.25 mm x 30 m, 0.25 μm film, Supelco, Bellefonte, PA). Helium was used as a carrier gas. The oven temperature was programmed from 160°C to 230°C at 4°C/min. Fatty acid methyl esters were identified by comparing the retention times to those of known standards. Inclusion of lipid standards with odd chain fatty acids permitted quantification of the amount of lipid in the sample. Dipentadecanoyl phosphatidylcholine (C15:0), diheptadecanoin (C17:0), trieicosenoin (C20:1), and cholesteryl eicosenoate (C20:1) were used as standards.

### Blood chemistry

Blood was collected following decapitation of the mice and collection of the blood via capillary using Microcuvette CB 300 K2E coated with K2 EDTA (16.444.100). All the blood collected in this study was sampled randomly (i.e. no fasting period) between 9 and 11 am to limit the effect of the circadian rhythm. Mice were subsequently dissected as soon as possible to limit the effect of fasting. Samples were then spun at 2000 g for 5 min at room temperature to extract plasma. Samples were pooled when large assay volume were required. Analysis of albumin, total protein, ALP, ALT, AST, bilirubin, iron, and NEFA (P19 only) were performed at the National Mouse Metabolic Phenotyping Center (MMPC) at the University of Massachusetts Medical School using a Cobas Clinical Chemistry Analyzer (Roche Diagnostics, Indianapolis, IN, USA) while plasma non-esterified fatty acid (NEFA) levels were measured photometrically using a kit (Zenbio, Durham, NC), according to the manufacturer’s protocol. Analysis of glucose, triglycerides and NEFA (P9-P13) was performed at Comparative Clinical Pathology Services, LLC., Columbia, Missouri, using commercially available assays on a Beckman-Coulter AU680 Automated Clinical Chemistry analyzer (Beckman-Coulter, Inc., Brea, CA). Triglyceride and glucose assays were obtained from Beckman-Coulter and the assay for non-essential Fatty Acids from Randox Laboratories (Randox Laboratories, Ltd., Kearneysville, West Virginia). In this study, we also used Luminex xMAP technology. The multiplexing analysis was performed using the Luminex™ 100 system (Luminex, Austin, TX, USA) by Eve Technologies Corp. (Calgary, Alberta). Eleven markers were simultaneously measured in the samples using a MILLIPLEX Mouse Cytokine/Chemokine 11-plex kit (Millipore, St. Charles, MO, USA) according to the manufacturer’s protocol. The 11-plex consisted of Amylin (active), C-Peptide 2, GIP (total), GLP-1 (active), ghrelin (active), glucagon, insulin, leptin, PP, PYY and Resistin. The assay sensitivities of these markers range from 1-23 pg/mL for the 11-plex. Individual analyte values are available in the MILLIPLEX protocol. IGF-1 was measured in the samples using a R&D Systems Mouse 1-Plex Luminex Assay (R&D Systems, Minneapolis, MN, USA) according to the manufacturer’s protocol. The assay sensitivity of this marker is 3.46 pg/mL. Adiponectin was measured in the samples using MILLIPLEX Mouse Cytokine/Chemokine 1-plex kit (Millipore, St. Charles, MO, USA) according to the manufacturer’s protocol. The assay sensitivity of this marker is 3 pg/mL. For experiments using Luminex system, if analytes were too low to be identified and outside of the dynamic range, it was deemed to be zero and reflected as such on dot plot graphs.

### Proteomic analysis

Proteomic analysis was performed in Dr. Parson’s laboratory and a work flow is presented in Supp Fig 5. Protein extraction, peptide tandem mass tagging and fractionation were performed by the FingerPrints Proteomics facilities at the University of Dundee. Protein samples were thawed, and proteins were extracted from each sample using Tris-HCl buffer (100 mM, pH 8.5) containing 4% SDS and 100 mM DTT. Samples are then processed using FASP protocol[64] with some modifications. After, removal of SDS with 8 M urea, proteins were alkylated with iodoacetamide and filters were washed 3 times with 100 mM Tris-HCL pH 8 then twice with 100 mM triethyl ammonium bicarbonate (TEAB). Proteins on the filters are then digested twice at 30°C with trypsin (2 x 2 μg), first overnight and then for another 6h in a final volume of 200 μl. Resulting tryptic peptides were desalted using C18 solid phase extraction cartridge (Empore, Agilent technologies) dried, dissolved in 100 mM TEAB and quantified using Pierce Quantitative Colorimetric Peptide Assay (Thermo Scientific). 100 μg of desalted tryptic peptides per sample were dissolved in 100 μl of 100 mM TEAB. The 10 different tandem mass tag labels comprising the TMT10plex™ kit (Thermo Fisher Scientific) were dissolved in 41 μL anhydrous acetonitrile. Each dissolved label was added to a different sample. Samples were labelled as follows: **Sample B** – Tag 127N Liver from 3 WT mice at P0; **Sample D** – Tag 128N Liver from 3 *Smn^2B/-^* mice at P0; **Sample G** – Tag 129C Liver from 3 WT mice at P2; **Sample I** – Tag 130C Liver from 3 *Smn^2B/-^* mice at P2 (this was part of a wider proteomic screen, hence the discontinuous lettering). The sample-label mixture was incubated for 1 hour at room temperature. Labelling reaction was stopped by adding 8 μl of 5% hydroxylamine per sample. Following labelling with TMT, samples were mixed, desalted, and dried in a speed-vac at 30°C. Samples were re-dissolved in 200 μl ammonium formate (NH_4_HCO_2_) (10 mM, pH 10) and peptides were fractionated using an Ultimate 3000 RP-High pH High Performance Liquid Chromatography column (Thermo-Scientific) containing an XBridge C18 column (XBridge peptide BEH, 130Å, 3.5 μm, 2.1 × 150 mm) (Waters, Ireland) with an XBridge guard column (XBridge, C18, 3.5 μm, 2.1 × 10 mm) (Waters, Ireland). Buffers A and B used for fractionation consist, respectively, of (A) 10 mM ammonium formate in milliQ water and (B) 10 mM ammonium formate with 90% acetonitrile. Before use, both buffers were adjusted to pH 10 with ammonia. Fractions were collected using a WPS-3000FC auto-sampler (Thermo-Scientific) at 1 minute intervals. Column and guard column were equilibrated with 2% Buffer B for twenty minutes at a constant flow rate of 0.2 ml/min. 175 μl per sample was loaded onto the column at a rate of 0.2 ml/min, and the separation gradient was started 1 minute after sample was loaded onto the column. Peptides were eluted from the column with a gradient of 2% Buffer B to 5% Buffer B in 6 minutes, and then from 5% Buffer B to 60% Buffer B in 50 minutes. Column was washed for 16 minutes in 100% Buffer B and equilibrated at 2% Buffer B for 20 minutes as mentioned previously. The fraction collection started 1 minute after injection and stopped after 80 minutes (total 80 fractions, 200 μl each). The total number of fractions concatenated was set to 15 and the content of the fractions was dried and suspended in 50 μl of 1% formic acid prior to analysis with LC-MS.

### LC-MS/MS Analysis

Liquid chromatography-tandem mass spectrometry was performed by FingerPrints Proteomics Facilities at the University of Dundee, to the following protocol: Analysis of peptide readout was performed on a Q Exactive™ HF Hybrid Quadrupole-Orbitrap™ Mass Spectrometer (Thermo Scientific) coupled with a Dionex Ultimate 3000 RS (Thermo Scientific). LC buffers were made up to the following: Buffer A (2% acetonitrile and 0.1% formic acid in Milli-Q water (v/v)) and Buffer B (80% acetonitrile and 0.08% formic acid in Milli-Q water (v/v). Aliquots of 15 μL per sample were loaded at a rate of 5 μL/min onto a trap column (100 μm × 2 cm, PepMap nanoViper C18 column, 5 μm, 100 Å, Thermo Scientific) which was equilibrated with 98% Buffer A. The trap column was washed for 6 minutes at the same flow rate and then the trap column was switched in-line with a resolving C18 column (Thermo Scientific) (75 μm × 50 cm, PepMap RSLC C18 column, 2 μm, 100 Å). Peptides were eluted from the column at a constant flow rate of 300 nL/min with a linear gradient from 95% Buffer A to 40% Buffer B in 122 min, and then to 98% Buffer B by 132 min. The resolving column was then washed with 95% Buffer B for 15 min and re-equilibrated in 98% Buffer A for 32 min. Q Exactive™ HF was used in data dependent mode. A scan cycle was comprised of a MS1 scan (m/z range from 335-1800, with a maximum ion injection time of 50 ms, a resolution of 120,000 and automatic gain control (AGC) value of 3×10^6^) followed by 15 sequential-dependent MS2 scans (with an isolation window set to 0.4 Da, resolution at 60,000, maximum ion injection time at 200 ms and AGC 1×10^5^. To ensure mass accuracy, the mass spectrometer was calibrated on the first day that the runs were performed.

### Database search and protein identifications

Raw MS data from the 15 fractions were searched against mouse (*Mus musculus*) protein sequences from UniProtKB/Swiss-Prot (Version 20160629) using the MASCOT search engine (Matrix Science, Version 2.4) through Proteome Discoverer™ software (Version 1.4.1.14, Thermo Fisher). Parameters for database search were as follows: MS1 Tolerance: 10ppm; MS2 Tolerance: 0.06da; fixed modification: Carbamidomethyl (C) Variable Modification: Oxidation (M), Dioxidation (M), Acetyl (N-term), Gln->pyro-Glu (N-term Q), TMT 10(N-term and K); maximum missed cleavage: 2; and target FDR 0.01. All identifications were quantified as relative ratios of expression compared to control (WT at P0) through Proteome Discoverer™ software (Thermo Fisher, Version detailed above). Relative ratios along with UnitProtKB/Swiss-Prot identifications were exported into Microsoft Excel as a raw data file for further in-silico analysis.

### In-Silico Analysis

Mass spec data (from above) was manually subdivided into four distinct groups - Group A (changed at P0 but not at P2), B (changed at P0 and P2), C (not changed at P0, but changed at P2) and NS, depending on the protein expression changes at P0 and P2 with level of significance identified as expression change increased or decreased by 20%. This procedure allows proteins most likely to be involved in the development of pathology, namely those altered at P0 and P2 or P2 only (Groups B and C) to be identified. These subgroups were then uploaded into the BioLayout *Express3D* for expression profile clustering, DAVID functional annotation for enrichment analysis or Ingenuity Pathway analysis (IPA) for hierarchical cascade mapping and upstream regulator prediction. See below.

### BioLayoutExpress3D

BioLayout*Express*3D[65] is a tool for visualization and clustering data. Routinely, proteomic data sets are uploaded to BioLayout *Express3D*, a Pearson’s correlation coefficient (*r*-value) is used to measure similarity between protein expression profiles and a threshold for the Pearson’s correlation coefficient is set. The data set is then visualized as nodes (proteins) that are connected to each other in a network based on their expression levels (edges). This data set can further be subdivided into discreet “clusters” based on a Markov Clustering Algorithm (MCL), thus segregating data in an unbiased manner as previously described[66–68].

### DAVID

The Database for Annotation, Visualization and Integrated Discovery (DAVID) provides a widely accepted set of functional annotation tools to interrogate the molecular composition of data sets relative to known findings in the current literature[69, 70]. The functional clustering tool divides a list of proteins into functional protein groups, each with a different Enrichment Score (ES), thus assigning a significance value. Analysis where appropriate was carried out as previously described[66, 68].

### Statistics

Data are presented as the mean ± standard error of the mean. A two-sided Student’s *t* test was performed using Microsoft Excel or Graphpad Prism 7 to compare the means of data when only two groups were compared (i.e. wild type vs. *Smn^2B/-^*). One-way ANOVA analysis and two-way ANOVA were also used to distinguish differences between more than two groups when multiple comparisons were necessary (i.e. wild type vs. *Smn^2B/+^* vs. *Smn^2B/-^*) or additional variables were present. The post-test used for the ANOVA was either Tukey or Sidak. Significance was set at *P* ≤ 0.05 for *, *P* ≤ 0.01 for **, *P* ≤ 0.001 for *** and *P* ≤ 0.0001 for ****. N number for each experiment is as indicated in the figure legends.

### Data and materials availability

All authors had access to the study data and had reviewed and approved the final manuscript. All data associated with this study are available in the main text or the supplementary materials. Raw data can be provided upon request.

## List of abbreviations

AGC: automatic gain control
ALP: alkaline phosphatase
ALT: alanine aminotransferase
AST: aspartate aminotransferase
Bax: BCL2 associated X protein
DAVID: The Database for Annotation, Visualization and Integrated Discovery
ES: Enrichment Score
FasR: Fas receptor
H&E: Hematoxylin & eosin
HFD: high fat diet
IGF-1: insulin-like growth factor 1
IGFbp1: insulin like growth factor binding protein 1
IGF1R: insulin like growth factor 1 receptor
igfals: insulin like growth factor binding protein acid labile subunit
IPA: ingenuity pathway analysis
MCD: methionine and choline deficient diet
MCL: Markov Clustering Algorithm
NAFLD: non-alcoholic fatty liver disease
NASH: non-alcoholic steatohepatitis
NEFA: non-esterified fatty acid
P: postnatal day
p21: cyclin dependent kinase inhibitor 1A
p53: tumor protein p53
PAS: Periodic acid-Schiff
SMA: spinal muscular atrophy
SMN1: Survival motor neuron 1
TMT: Tandem Mass Tagging
TNFR1: TNF receptor superfamily member 1A

## Acknowledgements

We would like to extend our gratitude to Eva Szunyogova, Sabrina Gagnon, My Tran Trung, and Rebecca Yaworski, the Vanderbilt Mouse Metabolic Phenotyping Center and the University of Massachusetts Medical School National Mouse Metabolic Phenotyping Center (MMPC) for assistance with experiments. We also thank Dr. Lyndsay Murray for providing some tissues for the present study.

