## Supplemental material for "A mouse model for spinal muscular atrophy provides insights into non-alcoholic fatty liver disease pathogenesis"

***Supplementary Materials***

*Supplementary Figures*

**Supplementary Fig 1** IPA analysis of group B identify metabolism but also cell cycle pathways.

**Supplementary Fig 2** *Smn*$^{2B/-}$ liver mitochondria function is not compromised even in the presence of fatty acids.

**Supplementary Fig 3** Major metabolic hormone levels are largely unchanged in the plasma of *Smn*$^{2B/-}$ mice.

**Supplementary Fig 4** Non-cropped raw western blot data.

**Supplementary Fig 5** Proteomic workflow for experimentation and analysis**.**

*Supplementary tables*

**Supplementary Table 1**. Primers used in this study

**Supplementary Table 2**. Oxygraph protocol in the absence of fatty acids

**Supplementary Table 3**. Oxygraph protocol in the presence of fatty acids

### ***Supplementary Figures***

A

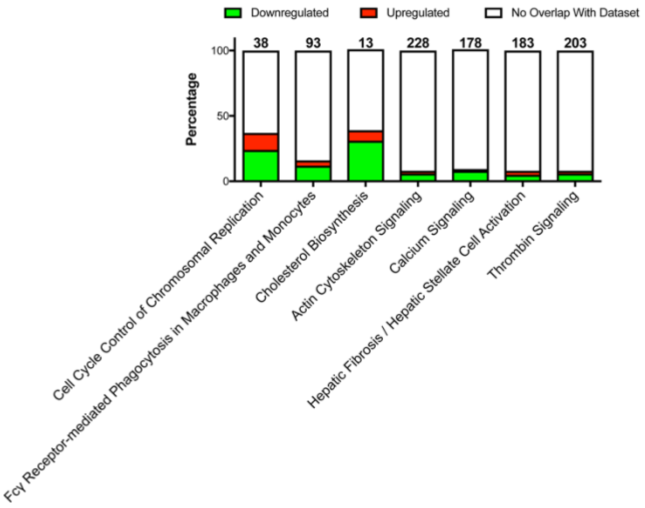

B

| Diseases & Functions | p-Value | # Molecules |
|---|---|---|
| Metabolism | 4,13E-10 | 49 |
| DNA Replication, Recombination, and Repair | 4,41E-10 | 36 |
| Cell Death | 4,7E-10 | 254 |
| Cellular Growth and Proliferation | 3,17E-08 | 135 |

**Supplementary Fig 1 IPA analysis of group B identify metabolism but also cell cycle pathways.** (A) IPA top canonical pathways highlighting the main disrupted cascades in Group B data set. Stacked bar chart displays the percentage of proteins that were upregulated (red), downregulated (green), and proteins that did not overlap with our data set (white) in each canonical pathway. The numerical value at the top of each bar represents the total number of proteins in the canonical pathway. (B) Top diseases and functions linked to our Group B data set identified by IPA functional analysis

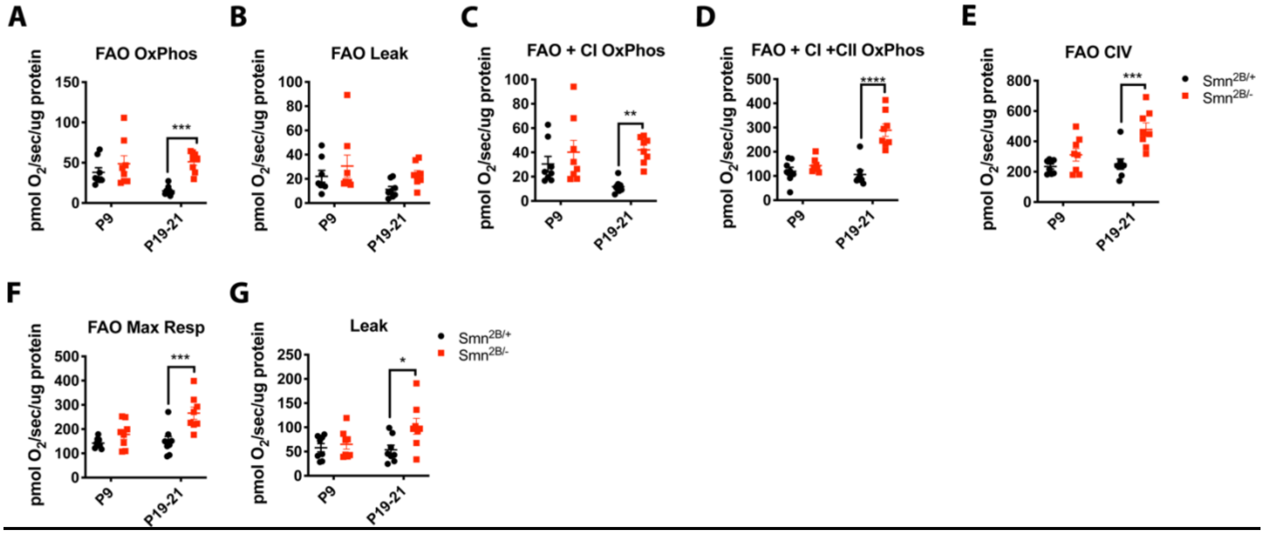

**Supplementary Fig 2 *Smn*$^{2B/-}$ liver mitochondria function is not compromised even in the presence of fatty acids.** High resolution respirometry of *Smn*$^{2B/-}$ hepatic mitochondria shows increased leak and higher respiration capacity at P19 but not at P9 in comparison to *Smn*$^{2B/+}$ hepatic mitochondria, suggesting intact mitochondrial function in the presence of fatty acids. (N value for each experiment is as follows: N = 8 for all experiments, two-way ANOVA with Sidak's multiple comparisons test, $P \leq 0.05$ for *, $P \leq 0.01$ for **, $P \leq 0.001$ for *** and $P \leq 0.0001$ for ****)

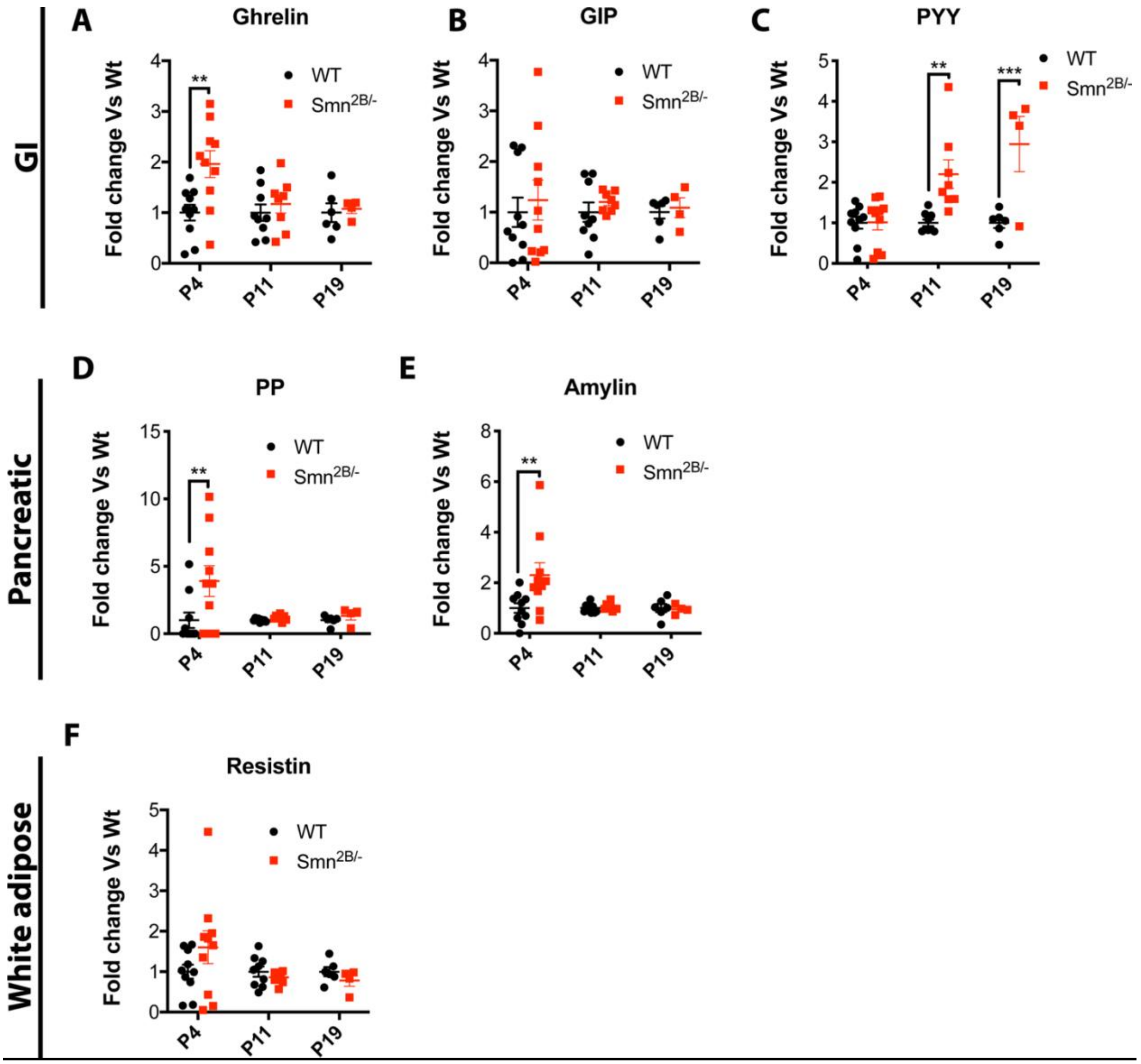

**Supplementary Fig 3 Major metabolic hormone levels are largely unchanged in the plasma of *Smn2B/-* mice.** (A-C) PYY is the only significantly changed hormone originating from the gastrointestinal system while ghrelin and GIP were largely unchanged. (D-E) Minor differences are present in pancreatic hormones. (H-J) No changes in resistin were observed were observed. (N value for each experiment is as follows: N = 8-10 for P4, P11 and 4-6 at P19 in A-F, two-way ANOVA with Sidak's multiple comparisons test, $P \leq 0.05$ for *, $P \leq 0.01$ for **)

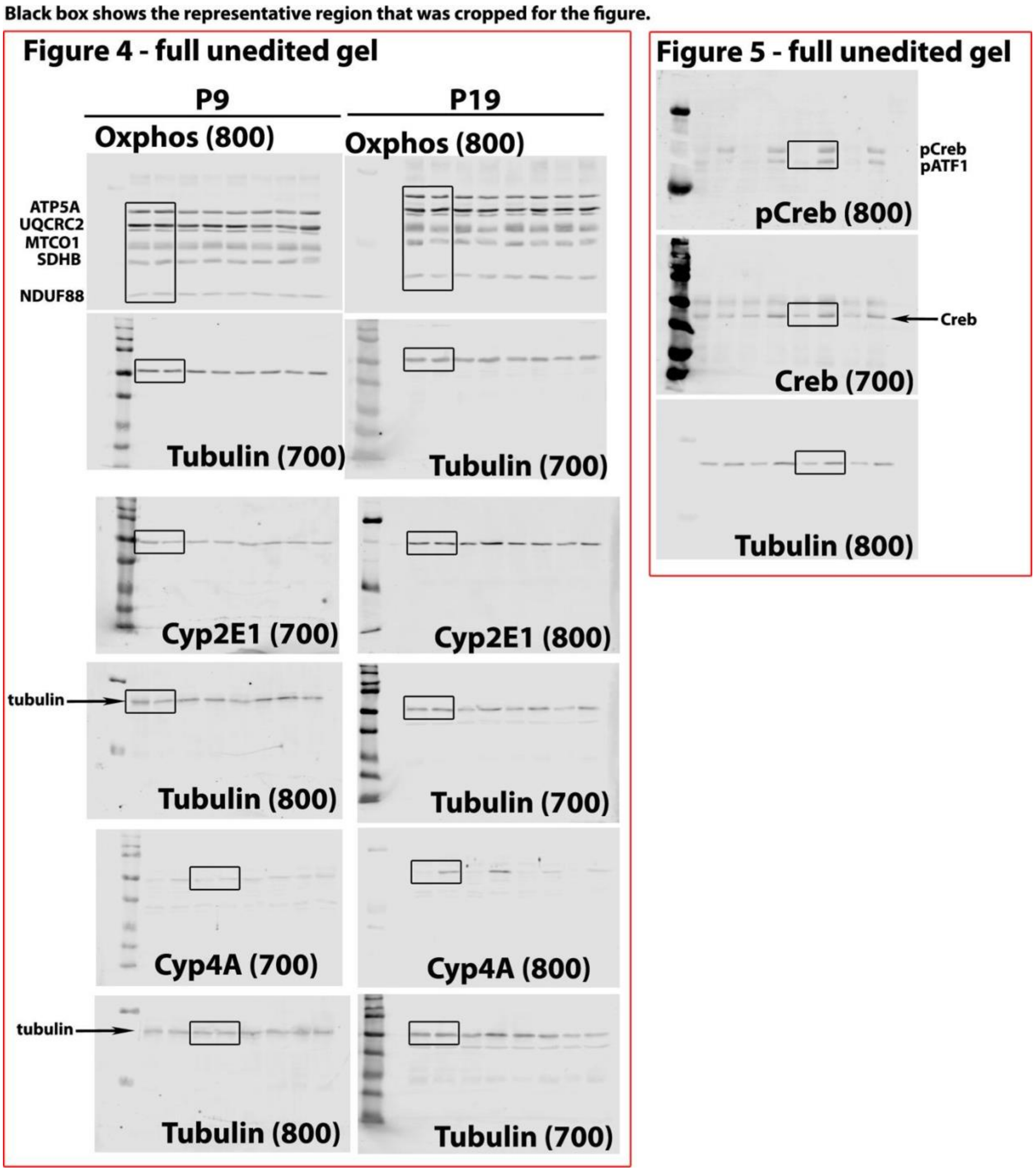

**Supplementary Fig 4 Non-cropped raw western blot data.** Raw western blots presented per Fig. Protein probed by antibody as well as channel used for identification via Odyssey acquisition system are written on each image. The numbers identify either the 700 or 800 nm wavelength fluorescence

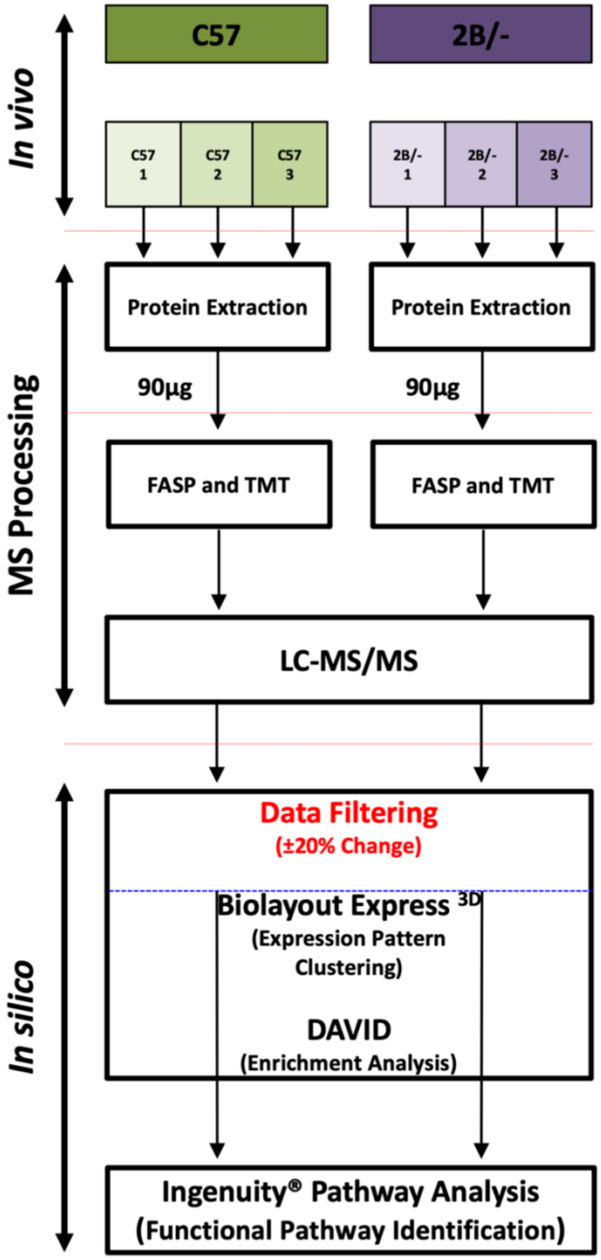

**Supplementary Fig 5 Proteomic workflow for experimentation and analysis.**

***Supplementary tables***

### Supplementary Table 1. Primers used in this study

| Gene name | Short form | Forward | Reverse | PrimePCR |
|---|---|---|---|---|
| **TNF Receptor Superfamily Member 6** | FasR | TGTGAACATGGAACCCTTGA | TTCAGGGTCATCCTGTCTCC | |
| **TNF Receptor Superfamily Member 1A** | TNFR1 | CCGGGAGAAGAGGGATAGCTT | TCGGACAGTCACTCACCAAGT | |
| **Caspase 8** | Casp8 | GGCCTCCATCTATGACCTGA | TGTGGTTCTGTTGCTCGAAG | |
| **BCL2 Associated X, Apoptosis Regulator** | Bax | TGCAGAGGATGATTGCTGAC | GATCAGCTCGGGCACTTTAG | |
| **BH3 interacting domain death agonist** | Bid | | | qMmuCID0022679 |
| **Tumor Protein P53** | p53 | GCTTCTCCGAAGACTGGATG | CTTCACTTGGGCCTTCAAAA | |
| **Cyclin-dependent kinase inhibitor 1A (P21)** | p21 | | | qMmuCED0046265 |
| **Transformed mouse 3T3 cell double minute 2** | Mdm2 | | | qMmuCID0025320 |
| **Complement C1r** | C1R | AACCATATTACAAGATGCTGACCA | CCTTGGGCTGTGCAGGTA | |
| **Complement C1s** | C1S | GGTGGATACTTCTGCTCCTGTC | AGGGCAGTGAACACATCTCC | |
| **Complement C1q B chain** | C1qb | CGTCGGCCCTAAGGGTACT | GGGGCTGTTGATGGTCCTC | |
| **Complement C3** | C3 | CCAGCTCCCCATTAGCTCTG | GCACTTGCCTCTTTAGGAAGTC | |
| **Complement C4** | C4 | TCTCACAAACCCCTCGACAT | AGCATCCTGGAACACCTGAA | |
| **Complement C5** | C5 | AGGGTACTTTGCCTGCTGAA | TGTGAAGGTGCTCTTGGATG | |
| **Complement C6** | C6 | | | qMmuCID0025195 |
| **Complement Factor B** | Factor B | GAGCGCAACTCCAGTGCTT | GAGGGACATAGGTACTCCAGG | |
| **Coagulation Factor II, Thrombin** | F2 | | | qMmuCED0046327 |
| **Coagulation Factor V** | F5 | CATGGAAACCTTACCGACAGAAA | CATGTGCCCCTTGGTATTGC | |
| **Coagulation Factor VII** | F7 | CGTCTGCTTCTGCCTCTTAGA | ATTTGCACAGATCAGCTGCTCAT | |
| **Coagulation Factor IX** | F9 | GCAAAACCGGGTCAAATCC | ACCTCCACAGAATGCCTCAATT | |
| **Coagulation Factor X** | F10 | | | qMmuCED0048020 |
| **Protein C, Inactivator Of Coagulation** | ProC | | | |

| | | | | |
|---|---|---|---|---|
| **Factors Va And VIIIa** | | | | |
| **Protein S** | ProS | | | qMmuCED0045958 |
| **Protein Z, Vitamin K Dependent Plasma Glycoprotein** | ProZ | | | |
| **Thrombopoietin** | Thpo | | | qMmuCED0037967 |
| **Hepatocyte Nuclear Factor 4 Alpha** | Hnf4a | AGAGGTTCTGTCCCAGCAGATC | CGTCTGTGATGTTGGCAATC | |
| **Insulin-like growth factor 1** | IGF1 | | | qMmuCID0005726 |
| **Insulin-like growth factor I receptor** | IGF1R | | | qMmuCID0005315 |
| **Insulin-like growth factor binding protein, acid labile subunit** | IGFals | | | qMmuCID0008201 |
| **Insulin-like growth factor binding protein 1** | IGFbp1 | | | qMmuCID0027402 |
| **Insulin-like growth factor binding protein 3** | IGFbp3 | | | qMmuCID0005232 |
| **Hepcidin** | Hamp | CCTATCTCCATCAACAGATG | AACAGATACCACACTGGGAA | |
| **Transferrin** | TF | CCATCCCATCACAACAAGGTATC | GCTAGTGTCCGATGCCTTCAC | |
| **Heme Oxygenase** | Hmox1 | GCCACCAAGGAGGTACACAT | GCTTGTTGCGCTCTATCTCC | |
| **Ceruloplasmin** | Cp | TCTACCAAGGAGTAGCCAGGA | ATCTTCCCTCTCATCCGTGC | |
| **Ferritin light chain** | L-Ferritin | CGTCTCCTCGAGTTTCAGAAC | CTCCTGGGTTTTACCCCATTC | |
| **Ferritin heavy chain** | H-Ferritin | CCATCAACCGCCAGATCAAC | GCCACATCATCTCGGTCAAA | |
| **Solute Carrier Family 40 Member 1** | Ferroportin | GCTGCTAGAATCGGTCTTTGGT | CAGCAACTGTGTCACCGTCAA | |
| **Solute carrier family 11 (proton-coupled divalent metal ion transporters), member 2** | Nramp2 | | | qMmuCID0016356 |
| **tyrosine 3-monooxygenase/ tryptophan 5-monooxygenase activation protein, zeta polypeptide** | Ywhaz | AAGACAGCACGCTAATAATGC | TTGGAAGGCCGGTTAATTTTC | |
| **succinate dehydrogenase** | Sdha | GCCTGGTCTGTATGCCTGTG | CCGATTCTTCTCCAGCATTTG | |

| | | | |
|---|---|---|---|
| **complex, subunit A, flavoprotein** | | | |
| **polymerase (RNA) II (DNA directed) polypeptide J** | Polr2j | ACCACACTCTGGGGAACATC | CTCGCTGATGAGGTCTGTGA |
| **hypoxanthine guanine phosphoribosyl transferase** | Hprt1 | CCCAGCGTCGTGATTAGTGATG | TTCAGTCCTGTCCATAATCAGTC |

**Supplementary Table 2. Oxygraph protocol in the absence of fatty acids**

| Substrate | Volume | Concentration of Stock | Concentration of substrate in the chamber |
|---|---|---|---|
| **Amplex ultra red** | 2 uL | 10 mM | 50 uM |
| **Horseradish peroxidase** | 10 uL | 10 mM | 10 U/mL |
| **H2O2** | 5 ul | 40 uM | 0.1 uM titrations X3 |
| **800 mM Malate** | 5 uL | 800 mM | 2 mM |
| **Pyruvate** | 10 uL | 2 M | 5 mM |
| **ADP/Mg2+** | 20 uL, 20 uL | 500 mM | 5 mM |
| **Glutamate** | 10 uL | 2 M | 10 mM |
| **Succinate** | 20 uL | 1 M | 10 mM |
| **500 mM ADP/Mg** | 20 uL, 20 uL | 500 mM | 5 mM |
| **Oligomycin** | 1 ul | 5 mM | 2.5 μM |
| **FCCP** | 0.5 uL titrations until max respiration | 1 mM | 0.25 uM titration |
| **Antimycin A** | 1 uL | 5 mM | 2.5 uM |
| **Ascorbate + TMPD** | 5 uL, 5 uL | 800 mM Asc, 200 mM TMPD | 2 mM Asc, 0.5 mM TMPD |
| **Sodium Azide** | 50 uL | 4 M | 100 mM |

**Supplementary Table 3. Oxygraph protocol in the presence of fatty acids**

| Substrate | Volume | Concentration of Stock | Concentration of substrate in the chamber |
|---|---|---|---|
| **800 mM Malate** | 5 uL | 800 mM | 2 mM |
| **Oct Car** | 4 uL | 100 mM | 0.2 mM |
| **ADP/Mg2+** | 20 uL, 20 uL | 500 mM | 5 mM |
| **Pyruvate** | 10 uL | 2 M | 5 mM |
| **Glutamate** | 10 uL | 2 M | 10 mM |
| **Succinate** | 20 uL | 1 M | 10 mM |
| **500 mM ADP/Mg** | 20 uL, 20 uL | 500 mM | 5 mM |
| **Oligomycin** | 1 ul | 5 mM | 2.5 uM |
| **FCCP** | 0.5 uL titrations until max respiration | 1 mM | 0.25uM titration |
| **Antimycin A** | 1 uL | 5 mM | 2.5 uM |
| **Ascorbate + TMPD** | 5 uL, 5 uL | 800 mM Asc, 200 mM TMPD | 2 mM Asc, 0.5 mM TMPD |
| **Sodium Azide** | 50 uL | 4 M | 100 mM |
